# A functional connectome phenotyping dataset including cognitive state and personality measures

**DOI:** 10.1101/164764

**Authors:** Natacha Mendes, Sabine Oligschläger, Mark E. Lauckner, Johannes Golchert, Julia M. Huntenburg, Marcel Falkiewicz, Melissa Ellamil, Sarah Krause, Blazej M. Baczkowski, Roberto Cozatl, Anastasia Osoianu, Deniz Kumral, Jared Pool, Laura Golz, Maria Dreyer, Philipp Haueis, Rebecca Jost, Yelyzaveta Kramarenko, Haakon Engen, Katharina Ohrnberger, Krzysztof J. Gorgolewski, Nicolas Farrugia, Anahit Babayan, Andrea Reiter, H. Lina Schaare, Janis Reinelt, Josefin Röbbig, Marie Uhlig, Miray Erbey, Michael Gaebler, Jonathan Smallwood, Arno Villringer, Daniel S. Margulies

## Abstract

The dataset enables exploration of higher-order cognitive faculties, self-generated mental experience, and personality features in relation to the intrinsic functional architecture of the brain. We provide multimodal magnetic resonance imaging (MRI) data and a broad set of state and trait phenotypic assessments: mind-wandering, personality traits, and cognitive abilities. Specifically, 194 healthy participants (between 20 and 75 years of age) filled out 31 questionnaires, performed 7 tasks, and reported 4 probes of in-scanner mind-wandering. The scanning session included four 15.5-min resting-state functional MRI runs using a multiband EPI sequence and a high-resolution structural scan using a 3D MP2RAGE sequence. This dataset constitutes one part of the MPI-Leipzig Mind-Brain-Body database.

## Background & Summary

Understanding the unique features of brain organization giving rise to distinct patterns of behavior, cognition, and mental experience remains one of the key research questions in the emerging field of human functional connectomics (Kelly et al., 2012). Functional connectivity has become a prominent method for investigating phenotypic differences across individuals (Vaidya & Gordon, 2013; Smith et al., 2015). However, there is ever greater need for validation of findings across independent datasets. The dataset presented here joins several others in contributing to this research agenda (e.g., Nooner et al., 2012; Holmes et al., 2015; Van Essen et al., 2013) and provides an additional resource for cross-site validation studies.

We acquired a wide range of self-reported personality measures as well as features of self-generated mental experience. In addition, a core magnetic resonance imaging (MRI) dataset—including one-hour of resting-state fMRI data—was acquired on 194 healthy participants. Questionnaires and behavioral measures were acquired over several follow-up sessions.

This dataset constitutes one part of the MPI-Leipzig Mind-Brain-Body database (manuscript forthcoming). It enables exploration of individual variance across cognitive and emotional phenotypes in relation to the brain. All MRI data were acquired on the same 3T Siemens Verio MRI scanner.

## Methods

### Participants

In total, datasets from 194 native German-speaking participants are included (94 female, mean age = 34 years, median age = 27, SD = 16 years; Figure 1). All participants were scanned on a 3 Tesla magnetic resonance imaging (MRI) scanner (Siemens Verio 3T) for the acquisition of one structural and four resting-state functional MRI scans. In addition, extensive questionnaire and task performance data were acquired from each participant. A subset of participants (N=109) were also included in the complementary data acquisition by Babayan et al. (manuscript forthcoming).

**Figure 1.**
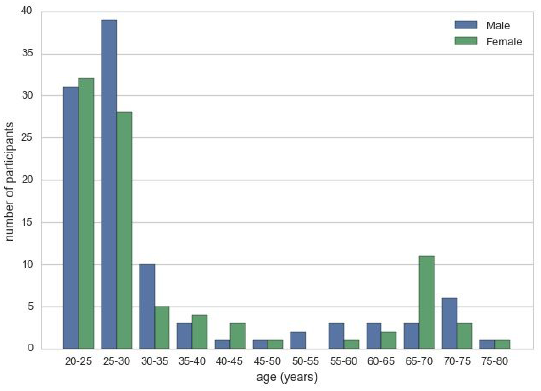
Age distribution. Age distribution (5-year bins) of the participants split by gender.

#### Recruitment and inclusion criteria

Prospective participants were initially recruited by the Leipzig Study for Mind-Body-Emotion Interactions project (manuscript forthcoming). Additional participants were recruited through online and poster advertisements. All participants were prescreened via telephone to determine their eligibility for the current study (Table 1). Participants fulfilling the eligibility criteria (including medical screening for MRI-scanning and neurological history) were invited to Max Planck Institute for Human Cognitive and Brain Sciences (MPI-CBS) where they were screened for past and present psychiatric disorders using the Structured Clinical Interview for DSM-IV (SCID-I; Wittchen et al., 1995). After meeting eligibility criteria, participants received detailed information regarding the study.

**Table 1.**
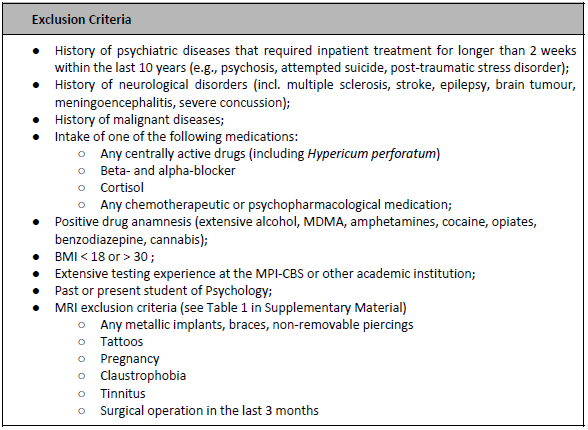
**Exclusion criteria.** Exclusion Criteria criteria to prospective participants.

All participants fulfilled the MRI safety requirements of the MPI-CBS (Supplementary Table 1), provided written informed consent (including agreement to their data being shared anonymously) prior to their participation in the study. Participants received monetary compensation for their participation. The study protocol was approved by the ethics committee of the University of Leipzig (097/15-ff).

#### Data acquisition and protocol overview

Participants were required to complete: 1) four functional MRI scans within one scanning session and, if not previously acquired in the study by Babayan et al (manuscript forthcoming), one structural scan; 2) a battery of cognitive, behavioral, and personality questionnaires spread over five appointments, and 3) a set of cognitive and creativity tasks spread over two appointments.

**Table 2.** **Phases of the data acquisition.** Overview of the different phases of the data acquisition.

| **Day 1**<br>**(MPI CBS)** | **Day 2**<br>**(Home)** | **Day 3**<br>**(Uni Clinic)** | **Day 4**<br>**(Home)** | **Day 5**<br>**(MPI CBS)** |
| --- | --- | --- | --- | --- |
| ASR | ACS | Scanning session | AMAS | BIS/BAS |
| God-MSI | BDI |  | NEO PI-R | CAQ |
| IAT | BP | FBI |  | MCQ-30 |
| IMIS | ESS | S-D-MW |  | BCQ |
| MGIQ | HADS | Short-NYC-Q_inscan1-4 |  | FFMQ |
| SCS | MMI | Short-NYC-Q_postETS |  | RAT |
| SD3 | MPU | NYC-Q_postscan |  | SYN |
| SE | PSSI | ETS |  | AUT |
| TPS |  | CCPT |  | TCIA |
| VISQ |  | Oddball |  |  |
| UPPS-P |  |  |  |  |
| SDS |  |  |  |  |

The data acquisition took place over five appointments over a two-year period (see Table 2):

- Day 1: We acquired data on a set of questionnaires that were completed at MPI-CBS (Tables 2 and 3).

**Table 3.** **Behavioral measures: questionnaires.** Overview of data available for each questionnaire.
- Day 2: We sent personalized links to participants, who could complete the set of online questionnaires at their convenience (Tables 2 and 3).
- Day 3: Participants were scanned at the Day Clinic for Cognitive Neurology, University of Leipzig. Before entering the scanner, participants completed a pen-and-paper practice trial of the short version of the New York Questionnaire (Ruby et al., 2013). While in the scanner, and immediately after each of the four resting state runs, participants received the computerized version of the same questionnaire. Immediately after the scanning session participants received additional questionnaires and a set of tasks (Tables 2, 3, and 4).

| Abbreviation | Behavioral Measure | N |
| --- | --- | --- |
| ACS | Attention Control Scale (Derryberry & Reed, 2002) | 210 |
| AMAS | Abbreviated Math Anxiety Scale (Hopko et al., 2003) | 145 |
| ASR | Adult Self Report (adapted from Achenbach & Rescorla, 2003) | 213 |
| BCQ | Body Consciousness questionnaire (Miller et al., 1981) | 79 |
| BDI | Beck Depressions Inventar-II (Beck et al., 1961, 1996; German version: Hautzinger et al., 1995) | 210 |
| BIS/BAS | Behavioral Inhibition and Approach System (Carver & White, 1994; Strobel et al., 2001; Gray, 1981, 1982) | 288 |
| BP | Boredom Proneness Scale (Farmer & Sundberg, 1986) | 209 |
| CAQ | Creative achievement questionnaire (Carson et al., 2005) | 79 |
| CCPT | Conjunctive continuous performance task (Shalev et al., 2011) | 169 |
| ESS | Epworth Sleepiness Scale (Johns, 1991; German version: Bloch et al., 1999) | 210 |
| FBI | Facebook Intensity Scale (adapted from Ellison et al., 2007) | 180 |
| FFMQ | Five facets of mindfulness (Baer et al., 2006) | 79 |
| Gold-MSI | Goldsmiths Musical Sophistication Index (German version: Schaal et al. 2014; Müllensiefen et al. 2014) | 214 |
| HADS | Hospital Anxiety and Depression Scale (Zigmond & Snaith, 1983; German version: Herrmann-Lingen, et al. 1995) | 210 |
| IAT | Internet Addiction Test (adapted from Young, 1998) | 214 |
| IMIS | Involuntary Musical Imagery Scale (adapted from Floridou et al., 2015) | 214 |
| MCQ-30 | Metacognition (Wells & Cartwright-Hatton, 2004; Sadeghi et al., 2014) | 79 |
| MGIQ | Multi-Gender Identity Questionnaire (adapted from Joel et al., 2014) | 159 |
| MMI | Multimedia Multitasking Index (Ophir et al., 2009) | 209 |
| MPU | Mobile Phone Usage (Developed in-house.) | 210 |
| NEO PI-R | NEO Personality Inventory-Revised (Ostendorf & Angleitner, 2004; Costa & McCrae, 1985, 1992) | 169 |
| NYC-Q_posttasks<br>NYC-Q_postscan<br>NYC-Q | New York Cognition Scale (Gorgolewski et al., 2014) | 202<br>188 |
| PSSI | Personality Style and Disorder Inventory (Kuhl & Kazén, 2009). | 209 |
| SCS | Brief Self-Control Scale (Tangney et al., 2004; German version adapted from: Bertrams & Dickhäuser, 2009) | 214 |
| SD3 | Short Dark Triad (adapted from Jones & Paulhus, 2014; Paulhus, 2013) | 214 |
| S-D-MW | Spontaneous and Deliberate Mind-Wandering (adapted from Carriere et al., 2013; see also Golchert et al. 2017) | 214 |
| SDS | Social Desirability Scale-17 (Crowne & Marlowe, 1960; German version: Stöber, 1999) | 214 |
| SE | Self-Esteem Scale (O'malley & Bachman, 1979) | 214 |
| Short-NYC-Q_inscan1<br>Short-NYC-Q_inscan2<br>Short-NYC-Q_inscan3<br>Short-NYC-Q_inscan4<br>Short-NYC-Q_postETS<br>Short-NYC-Q_prescan | Short New York Cognition Scale (Ruby et al., 2013) | 175<br>174<br>174<br>170<br>181<br>159 |
| TPS | Tuckman Procrastination Scale (Stöber, 1995; Tuckman, 1991). | 214 |
| UPPS-P | UPPS-P Impulsive Behavior Scale (Whiteside & Lynam, 2001; Lynam et al., 2006; see also Golchert et al., 2017) | 214 |
| VISQ | Varieties of Inner Speech (McCarthy-Jones & Fernyhough, 2011) | 214 |

**Table 4.** **Behavioral measures: tasks.** Overview of data available for each task.
- Day 4: The Abbreviated Math Anxiety Scale and the NEO Personality Inventory-Revised were completed online at the participant’s convenience.
- Day 5: We acquired data on a set of questionnaires and tasks that were administered at MPI-CBS. Tasks were conducted using pen-and-paper, computer-administered, as well as Limesurvey interfaces (Tables 2, 3, and 4).

| Abbreviation | Behavioral Measure | N |
| --- | --- | --- |
| AUT | Alternative uses task (Silvia, 2008; Guilford et al. 1978) | 77 |
| CCPT | Conjunctive continuous performance task (Shalev et al., 2011) | 169 |
| ETS | Emotional task switching (adapted from Whitmer & Banich, 2007; see Hildebrandt et al., 2016) | 189 |
| Oddball | Adaptive visual and auditory oddball target detection task (modified from: Huettel & McCarthy, 2004) | 137 |
| RAT | Remote associates test (Landmann et al., 2014; see also Lee et al., 2014) | 77 |
| SYN | Synesthesia color picker test (synesthete.org; Eagleman et al., 2007) | 73 |
| TCIA | Test of creative imagery abilities (Jankowska & Karwowski, 2015) | 77 |

Within each set of questionnaires and tasks, the order of presentation of questionnaires and tasks was randomized across participants. If participants failed to complete a given questionnaire it was excluded from data analysis. Due to dropout, not all participants completed the full set of questionnaires and tasks.

### Drug screening prior to MRI data acquisition

Each of the participants was instructed not to use illicit drugs within two weeks of the scanning appointment. Participants were also requested to abstain from alcohol and caffeine consumption, as well as nicotine on the night prior to the scanning day and on the day of scanning. Before the beginning of the MRI session, participants’ urine was biochemically screened with a MULTI 8/2 strip test (Diagnostik Nord, Schwerin, Germany) for the presence of buprenorphine (cutoff 10ng/mL), amphetamine (cutoff 1000ng/mL), benzodiazepine (300ng/mL), cocaine (cutoff 300ng/mL), methamphetamine (1000ng/mL), morphine/heroine (cutoff 300ng/mL), methadone (cutoff 300ng/mL), THC (cutoff 50ng/mL). Cutoff levels are those recommended by the American National Institute on Drug Abuse (NIDA; Hawks & Chiang, 1986). Participants provided informed consent on the use of the urine strip test and agreed to its anonymous data sharing, prior to their participation in the study.

### MRI data acquisition

All magnetic resonance imaging (MRI) data was acquired using a whole-body 3 Tesla scanner (Magnetom Verio, Siemens Healthcare, Erlangen, Germany) equipped with a 32-channel Siemens head coil at the Day Clinic for Cognitive Neurology, University of Leipzig. For each participant the following scans were obtained: 1) a high-resolution structural scan, 2) four resting-state functional MRI (rs-fMRI) scans, 3) two gradient echo fieldmaps and, 4) two pairs of spin echo images with reversed phase encoding direction. A low-resolution structural image of each participant was acquired using a FLAIR sequence for clinical screening.

#### Structural scan

The high-resolution structural image was acquired using a 3D MP2RAGE sequence (Marques et al., 2010) with the following parameters: voxel size = 1.0 mm isotropic, FOV = 256 ×240 ×176 mm, TR = 5000 ms, TE = 2.92 ms, TI1 = 700 ms, TI2 = 2500 ms, flip angle 1 = 4°, flip angle 2 = 5°, bandwidth = 240 Hz/Px, GRAPPA acceleration with iPAT factor 3 (32 reference lines), pre-scan normalization, duration = 8.22 min. From the two images produced by the MP2RAGE sequence at different inversion times (inv1 and inv2), a quantitative T1 map (t1map), and a uniform T1-weighted image (t1w) were generated. Importantly, the latter image is purely T1-weighted, whereas standard T1-weighted image, for example acquired with the MPRAGE sequence, also contain contributions of proton density and T2^*^. It should be taken into account that such differences can affect morphometric measures (Lorio et al., 2016).

For one participant, the structural scan is MPRAGE instead of MP2RAGE (the T1-weighted image file names contain the sequence type) with voxel size = 1 mm isotropic, FoV = 256 ×240 ×176, TR = 2300 ms, TE = 2.98 ms, TI = 900 ms, flip angle = 9°, bandwidth = 238 Hz/Px.

#### Resting-state scans

Four rs-fMRI scans were acquired in axial orientation using T2^*^-weighted gradient-echo echo planar imaging (GE-EPI) with multiband acceleration, sensitive to blood oxygen level-dependent (BOLD) contrast (Feinberg et al., 2010; Moeller et al., 2010). Sequences were identical across the four runs, with the exception of alternating slice orientation and phase-encoding direction, to vary the spatial distribution of distortions and signal loss. Thus, the y-axis was aligned parallel to the AC-PC axis for runs 1 and 2, and parallel to orbitofrontal cortex for runs 2 and 4. The phase-encoding direction was A–P for runs 1 and 3, and P–A for runs 2 and 4. Further parameters were set as follows for all four runs: voxel size = 2.3 mm isotropic, FOV = 202 ×202 mm^2^, imaging matrix = 88 ×88, 64 slices with 2.3 mm thickness, TR = 1400 ms, TE = 39.4 ms, flip angle = 69°, echo spacing = 0.67 ms, bandwidth = 1776 Hz/Px, partial fourier 7/8, no pre-scan normalization, multiband acceleration factor = 4, 657 volumes, duration = 15 min 30 s. During the resting-state scans, participants were instructed to remain awake with their eyes open and to fixate on a crosshair.

#### Scans for distortion correction

Two prominent methods exist to correct for geometric distortions in EPI images: fieldmaps, which represent the degree of distortion as calculated from two phase images with different echo times (Jezzard & Balaban 1995; Reber et al., 1998), and reverse phase encoding, in which pairs of “blip-up blip-down” images are acquired with opposite phase encoding direction — thus opposite distortions — and used to model a middle distortion-free image (Chang & Fitzpatrick, 1992; Andersson et al., 2003). This datasets contains scans required for both methods to accommodate different preprocessing approaches and facilitate method comparison. Before each pair of resting-state runs with the same y-axis orientation (see above), the following scans were acquired in the same orientation as the subsequent resting-state scans: a pair of spin echo images (voxel size = 2.3 mm isotropic, FOV = 202 ×202 mm^2^, imaging matrix = 88 ×88, 64 slices with 2.3 mm thickness, TR = 2200 ms, TE = 52 ms, flip angle = 90°, echo spacing = 0.67 ms, phase encoding = AP / PA, bandwidth = 1776 Hz/Px, partial fourier 6/8, no pre-scan normalization, duration = 0.20 min each), and a gradient echo fieldmap (voxel size = 2.3 mm isotropic, FOV = 202 ×202 mm^2^, imaging matrix = 88 ×88, 64 slices with 2.3 mm thickness, TR = 680 ms, TE1 = 5.19 ms, TE2 = 7.65 ms, flip angle = 60°, bandwidth = 389 Hz/Px, prescan normalization, no partial fourier, duration = 2.03 min).

#### Additional scans

109 subjects also took part in the protocol by Babayan et al. (manuscript forthcoming). Therefore, additional modalities might be available for these subjects. Modalities include high-resolution T2-weighted (108 subjects), diffusion-weighted (109), 3D FLAIR (47), phases and magnitudes of gradient-echo images suitable for Susceptibility-Weighted Imaging (SWI) and Quantitative Susceptibility Mapping (QSM) (45 subjects), as well as an additional 15-minute resting-state scan for all 109 subjects.

#### MRI data preprocessing

To enhance data usability we provide preprocessed data from 189 subjects.^1^ Data from five participants were further excluded due to failure at the preprocessing stage. The raw MRI data of these subjects are not corrupted, and are therefore available in the main database. Preprocessing pipelines were implemented using Nipype (Gorgolewski et al., 2011) and are described in more detail below. All code is openly available.^2^

Importantly, the preprocessing performed here is just one out of a multitude of possible pipelines that could be conceived for this dataset. The decisions taken at individual processing steps will not be suitable for every application. Users are strongly advised to familiarize themselves with the details of the workflow before adopting the preprocessed data for their study. We also encourage users to subscribe to the mailing list for updates and discussions regarding the preprocessing pipelines used here.^3^

#### Structural data

The background of the uniform T1-weighted image was removed using CBS Tools (Bazin et al., 2014), and the masked image was used for cortical surface reconstruction using FreeSurfer’s full version of recon-all (Dale et al., 1999; Fischl et al., 1999). A brain mask was created based on the FreeSurfer segmentation results. Diffeomorphic nonlinear registration as implemented in ANTs SyN algorithm (Avants et al., 2011) was used to compute a spatial transformation between the individual’s T1-weighted image and the MNI152 1mm standard space.

To remove identifying information from the structural MRI scans, a mask for defacing was created from the MP2RAGE images using CBS Tools (Bazin et al. 2014). This mask was subsequently applied to all anatomical scans.

#### Functional data

The first five volumes of each resting-state run were excluded. Transformation parameters for motion correction were obtained by rigid-body realignment to the first volume of the shortened time series using FSL MCFLIRT (Jenkinson et al., 2002). The fieldmap images were preprocessed using the fsl_prepare_fieldmap script. A temporal mean image of the realigned time series was rigidly registered to the fieldmap magnitude image using FSL FLIRT (Jenkinson & Smith 2001) and unwarped using FSL FUGUE (Jenkinson et al. 2012) to estimate transformation parameters for distortion correction. The unwarped temporal mean was rigidly coregistered to the subject’s structural scan using FreeSurfer’s boundary-based registration algorithm (Greve & Fischl 2009), yielding transformation parameters for coregistration. The spatial transformations from motion correction, distortion correction, and coregistration were then combined and applied to each volume of the original time series in a single interpolation step. The time series were masked using the brain mask created from the structural image (see above). The six motion parameters and their first derivatives were included as nuisance regressors in a general linear model (GLM), along with regressors representing outliers as identified by Nipype’s rapidart algorithm^4^, as well as linear and quadratic trends. To remove physiological noise from the residual time series, we followed the aCompCor approach as described by Behzadi et al. (2007). Masks of the white matter and cerebrospinal fluid were created by applying FSL FAST (Zhang et al., 2001) to the T1-weighted image, thresholding the resulting probability images at 99%, eroding by one voxel and combining them to a single mask. Of the signal of all voxels included in this mask, the first six principal components were included as additional regressors in a second GLM, run on the residual time series from the first GLM. The denoised time series were temporally filtered to a frequency range between 0.01 and 0.1 Hz using FSL, mean centered and variance normalized using Nitime (Rokem et al., 2009). The fully preprocessed time series of all for runs were temporally concatenated. To facilitate analysis in standard space, the previously derived transformation was used to project the full-length time series into MNI152 2mm space. The preprocessed data are made available in the subjects’ native structural space and MNI standard space, along with the subject’s brain mask and all regressors used for denoising.

### Data security and data anonymization procedures

Data for all participants was stored on our instance of the eXtensible Neuroimaging Archive Toolkit (XNAT, Marcus et al. 2007) v.1.6.5. at the MPI-CBS. Access to the initial project was restricted (via XNAT’s private project mode) to members of the Neuroanatomy & Connectivity Group at MPI-CBS for initial curation and quality assessment of data. All data comprised in the MPI-Leipzig Mind-Brain-Body database were derived from MPI-CBS so data import into XNAT was done from a local secured network.

A specially customized XNAT uploader was used to upload all participants’ data to XNAT. The native DICOM format was used for MRI data, whilst a standard ASCII (*.csv, *.txt) format was employed to upload all other experimental data such as surveys, test batteries, and demographical data.

The anonymization measures applied to the MRI data consisted of removal of DICOM header tags containing information which could lead to the identification of test subjects as well as the defacing of all structural (NIFTI) scans. Specific surveys and test batteries containing sensitive information are only available via the restricted project in XNAT for which access needs to be applied for (see the Usage Notes section below).

### Code availability

All code that was implemented for data acquisition and processing is available online.^5^ Data handling and computation of summary measures were implemented in Python. The pipeline used for MRI preprocessing is also available.^6^

The tasks that the participants received were implemented using the Python package PsychoPy2 Experiment Builder v1.81.03 (Peirce, 2007, 2008), OpenSesame 0.27.4 (Mathôt & Theeuwes, 2012) and Presentation^®^ software (Version 16.5, Neurobehavioral Systems, Inc., Berkeley, CA, <u>www.neurobs.com</u>). We provide the respective source codes of the adaptive visual and auditory oddball task (oddball) ^7^, conjunctive continuous performance task (CCPT) ^8^, and emotional task switching task (ETS)^7^.

### Data Records

#### Survey and task data

A comprehensive list of behavioral and questionnaire data are given in Supplementary Table 2. Data from all questionnaires are released as summary scores. Results of questionnaires without summary scores are released as raw item scores, namely: Multi-Gender Identity (MGIQ), Mobile phone usage (MPU), FBI (Facebook intensity scale), New York cognition (NYC-Q), and the short version of the New York cognition (Short-NYC-Q) questionnaires. Task data for the CCPT, ETS, and oddball task are available via subject-specific csv files. Accompanying specifications and information for each questionnaire and task are given in txt file format.

A basic demographic summary is provided together with general information on data acquisition. The metafile includes gender, age (5-year bins), current or past diagnosed psychiatric disorder(s), result of the drug test on day of scanning, and formal education.

#### MRI data

The dataset is organized in concordance with the Brain Imaging Data Structure (BIDS) format (Gorgolewski et al., 2016a). This facilitates data analysis, for example with BIDS-Apps (Gorgolewski et al., 2016b).^9^ BIDS-Apps encapsulate standard MRI analysis tools within an application that understands the BIDS format and allows to automatically access relevant data and metadata.

MRI data are currently available from two locations:

1. OpenfMRI.org platform also hosts the raw data.^10^
2. Gesellschaft für wissenschaftliche Datenverarbeitung mbH Göttingen (GWDG).^11^ The data at this location is accessible through web browser and fast FTP connection. It contains both raw and preprocessed data. Currently, preprocessed resting-state fMRI data is available.^12^ In the case the location of the data changes in the future, the location of the dataset can be resolved with PID 21.11101/0000-0004-2CD6-A.^13^

### Technical Validation

All datasets were manually assessed for missing or corrupt data. Further quality control of the data was applied to the MRI and behavioral measures, as described below.

### MRI data quality assessment

Preprocessed MRI data were assessed for quality using the mriqc package^14^ (Esteban et al. 2017), implemented in Python. mriqc creates a report for each individual scan based on assessment of movement parameters, coregistration, and temporal signal-to-noise (tSNR) calculations. For comparison, all individual-level scores are displayed with respect to the group-level distribution. We visually inspected the quality assessment reports for each subject to ensure adequate coregistration and fieldmap correction.

As motion during the resting-state fMRI scan poses a substantial source of noise (Power et al., 2014), we characterized motion for each run as the mean and maximum framewise displacement (Figure 2). Overall, the summary of motion parameters demonstrates that the data are largely of sufficient quality, with 89.2% of runs showing less than one voxel (2.3 mm) maximum framewise displacement, and a mean framewise displacement of 0.18 mm (SD = 0.08 mm).

**Figure 2.**
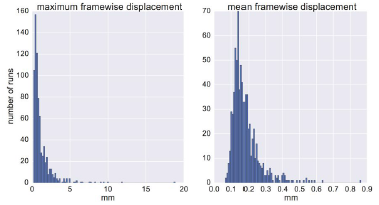
Quality assessment of resting-state fMRI scans. Distribution of motion (maximum and mean framewise displacement).

Fieldmap correction provides an approach to correct for distortions due to susceptibility artifacts. While unable to recover signal loss, the correction of such nonlinear distortions improves coregistration between scan types, and group-level alignment (Jezzard, 2012). As an example, we present a single dataset, pre- and post-fieldmap correction, in Figure 3. As expected, fieldmap correction primarily shifted voxels within ventral regions.

**Figure 3.**
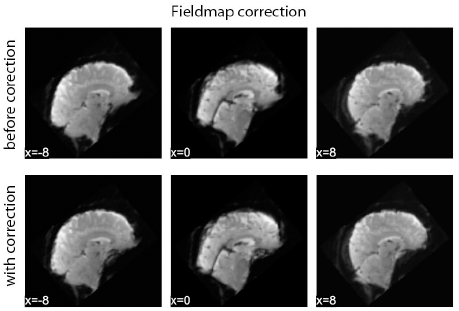
Example impact of fieldmap correction.

Temporal signal-to-noise (tSNR), which is calculated on the voxel-level as the mean signal divided by the standard deviation, offers a general overview of the local differences across the brain. We observed lower tSNR in ventral regions, including the orbitofrontal and temporal cortex (Figure 4).

**Figure 4.**
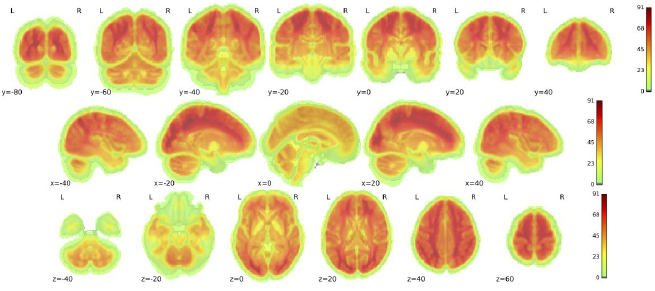
Temporal Signal-to-Noise (tSNR). Group-level variance in temporal signal-to-noise (tSNR) across the brain. tSNR values are lower in ventral regions including orbitofrontal and temporal cortex.

### Behavioral measures quality assessment

Fifteen questionnaires without a published German version were in-house translated (English-German). To ensure general usability of the translated questionnaires, their reliability was estimated using Cronbach’s Alpha coefficient (see Table 5). For comparison, the Cronbach’s Alpha coefficients from the original questionnaires are also reported in Table 5. Internal consistency of the majority of questionnaires was acceptable, with an average Cronbach’s Alpha of 0.78, thus showing that the German translations of those specific questionnaires are reproducible and valid. However, three questionnaires (Short Dark Triad, Body Consciousness questionnaires, and the Creative Achievement questionnaire) and four scales (two scales of the Five Facets of Mindfulness questionnaire, one scale of the Metacognition questionnaire, and one scale of the Involuntary Musical Imagery scale) showed modest reliability, with Cronbach’s Alpha coefficient < 0.70, and should be interpreted with caution.

**Table 5.** **Reliability of translated questionnaires.** Estimated reliability of the English-German translated questionnaires using Cronbach’s Alpha coefficient (α).

| Translated Questionnaires | Cronbach's alpha coefficient ($\alpha$) |
| --- | --- |
| Abbreviated math anxiety scale (Hopko et al., 2003) | $\alpha = .92$ (English original: $\alpha = .90$) |
| Attention control scale (Derryberry and Reed, 2002) | $\alpha = .74$ (English original: not reported) |
| Body consciousness questionnaire<br>(Miller and al., 1981) | *Private body scale, $\alpha = .63$*<br>*Public body scale, $\alpha = .62$*<br>*Body competence scale, $\alpha = .62$*<br>Note. Cronbach's $\alpha$ coefficients of the original scales are not available |
| Boredom proneness scale (Farmer and Sundberg, 1986) | $\alpha = .84$ (English original: $\alpha = .79$) |
| Creative achievement questionnaire<br>(Carson & Higgins, 2005) | $\alpha = .67$ (English original: $\alpha = .96$ ) |
| Five facets of mindfulness<br>(Baer et al., 2006) | *Observing scale*, $\alpha = .68$ (English original: $\alpha = .83$)<br>*Describing scale*, $\alpha = .89$ (English original: $\alpha = .91$)<br>*Acting with Awareness scale*, $\alpha = .70$ (English original: $\alpha = .87$)<br>*Nonjudging scale*, $\alpha = .87$ (English original: $\alpha = .87$)<br>*Nonreactivity scale*, $\alpha = .69$ (English original: $\alpha = .75$) |
| Internet addiction test (Young, 1999) | $\alpha = .91$ (item 3 was excluded from the analysis due to different scaling; values for English unavailable) |
| Involuntary musical imagery scale<br>(Floridou et al., 2015) | *Negative valence*, $\alpha = .88$ for (English original: $\alpha = .91$)<br>*Movement*, $\alpha = .92$ (English original: $\alpha = .88$)<br>*Personal reflections*, $\alpha = .64$ (English original: $\alpha = .76$)<br>*Help*, $\alpha = .90$ (English original: $\alpha = .84$) |
| Media-multitasking inventory<br>(Ophir et al., 2009) | $\alpha = .97$ (English original: not reported) |
| Metacognition<br>(Wells and Cartwright-Hatton, 2004) | *Cognitive confidence*, $\alpha = .80$ (English original: $\alpha = 0.93$)<br>*Positive beliefs*, $\alpha = .85$ (English original: $\alpha = 0.92$)<br>*Cognitive self-consciousness*, $\alpha = .85$ (English original: $\alpha = 0.92$)<br>*Uncontrollability and danger*, $\alpha = .80$ (English original: $\alpha = 0.91$)<br>*Need to control thoughts*, $\alpha = .67$ (English original: $\alpha = 0.72$) |
| Self-esteem scale (O'malley and Bachman, 1979) | $\alpha = .88$ (English original: $\alpha = .79$ for males; $\alpha = .83$ for females) |
| Short dark triad<br>(Paulhus, 2013; Original by Jones and Paulhus, 2014) | *Machiavellianism*, $\alpha = .68$ (English original: $\alpha = .78$)<br>*Narcissism*, $\alpha = .65$ (English original: $\alpha = .77$)<br>*Psychopathy*, $\alpha = .59$ for (English original: $\alpha = .80$) |
| Spontaneous and deliberate mind-wandering | *Deliberate mind wandering*, $\alpha = .81$ (English original: $\alpha = .90$) |
| (Carriere et al., 2013) | *Spontaneous mind wandering*, $\alpha = .81$ (English original: $\alpha = .88$) |
| UPPS-P impulsive behavior scale<br>(Whiteside and Lynam, 2001;<br>Lynam et al., 2006) | *Negative urgency*, $\alpha = .83$ (English original: $\alpha = .90$)<br>*Lack of premeditation*, $\alpha = .75$ (English original: $\alpha = .91$)<br>*Lack of perseverance*, $\alpha = .84$ (English original: $\alpha = .82$)<br>*Sensation seeking*, $\alpha = .82$ (English original: $\alpha = .86$)<br>*Positive urgency*, $\alpha = .90$ (English original: not reported) |
| Varieties of inner speech<br>(McCarthy-Jones and Fernyhough, 2011) | *Dialogic inner speech*, $\alpha = .74$ (English original: $\alpha = .83$)<br>*Condensed inner speech*, $\alpha = .79$ (English original: $\alpha = .83$)<br>*Other people in inner speech*, $\alpha = .86$ (English original: $\alpha = .88$)<br>*Evaluative inner speech*, $\alpha = .74$ (English original: $\alpha = .80$) |

### Usage Notes

The MRI dataset can be accessed at www.openfmri.org, and the behavioral data is available at www.nitrc.org^15^. The following data are publicly available: 1) MRI data (structural and functional), 2) general demographic of the studied population, 3) summary scores and/or indexes of the questionnaires and tasks, and 4) raw scores of the measures that do not possess summary scores and have not been classified as sensitive. All MRI datasets are made available in NIFTI format, and all anatomical scans have been defaced.

The dataset, protocols, and software used in the acquisition and processing of the data are documented, curated, and available for research purposes. For access to the behavioral data, users must first agree to the terms of data usage.

#### Additional access to sensitive behavioral measures

Individual behavioral scores and sensitive phenotypic measures may be made available upon request.^16^ The completion of additional data license and confidentiality forms will be required in advance of further data access.

## Acknowledgements

This work was partially supported by the Volkswagen Foundation (AZ.: 89 440). We thank Shameem Wagner and Elizabeth Kelly for assistance in the preparation of the manuscript.

## Author contributions

Conception, design, and preparation of the manuscript: D.S.M., J.G., J.M.H., M.E.L., M.F., N.M., S.O.

Behavioral data analyses: J.G., M.E.L., S.O.

MRI data preprocessing: J.M.H., M.E.L.

Quality Control of MRI data: D.S.M., J.G., J.M.H., M.E.L., S.O.

Contributions to study design: B.M.B., H.E., J.P., J.S., K.J.G., K.O., N.F.

Participant recruitment: A.O., J.G., N.M., M.E.L., P.H., R.J., S.K., Y.K.

Data acquisition: D.K., J.G., J.P., L.G., M.D., M.E.L., N.M., S.K., S.O.

Databasing: R.C.

Data contributions: A.B., A.R., A.V., H.L.S., J.R., M.E., M.G., M.U.

All authors provided critical feedback and approval of the manuscript.

## Competing interests

The authors declare no competing financial interests.

## Footnotes

1 Five participants did not have all four resting-state scans are available, and were excluded from preprocessing.

2 https://github.com/NeuroanatomyAndConnectivity/pipelines/tree/master/src/lsd_lemon

3 http://groups.google.com/group/resting_state_preprocessing

4 http://nipy.org/nipype/interfaces/generated/nipype.algorithms.rapidart.html

5 https://neuroanatomyandconnectivity.github.io/opendata/

6 https://github.com/NeuroanatomyAndConnectivity/pipelines/tree/v2.0/src/lsd_lemon (release v2.0)

7 https://github.com/NeuroanatomyAndConnectivity/opendata/tree/master/scripts

8 https://github.com/NeuroanatomyAndConnectivity/ConjunctiveContinuousPerformanceTask

9 http://bids-apps.neuroimaging.io

10 https://www.openfmri.org/dataset/ds000221/

11 https://www.gwdg.de/

12 The current URL for dataset is <u>ftp://ftp.gwdg.de/pub/misc/MPI-Leipzig_Mind-Brain-Body/</u> or https://ftp.gwdg.de/pub/misc/MPI-Leipzig_Mind-Brain-Body/

13 e.g., https://hdl.handle.net/21.11101/0000-0004-2CD6-A

14 The code was adapted from https://github.com/chrisfilo/mriqc and can be found at https://github.com/NeuroanatomyAndConnectivity/pipelines/tree/master/src/lsd_lemon (release v2.0)

15 http://nitrc.org/projects/mpilmbb/

16 Contact corresponding authors.

## References

Achenbach, T. M. & Rescorla, L. A. in (Research Center for Children, Youth, & Families, University of Vermont, Burlington, VT, USA, 2003).

Andersson, J. L., Skare, S. & Ashburner, J. How to correct susceptibility distortions in spin-echo echo-planar images: application to diffusion tensor imaging. Neuroimage 20, 870–888 (2003).

Avants, B. B. et al. A reproducible evaluation of ANTs similarity metric performance in brain image registration. Neuroimage 54, 2033–2044 (2011).

Babayan et al. A dataset to investigate mind-brain-body interactions in younger and older adults with a focus on emotions. (Forthcoming).

Baer, R. A., Smith, G. T., Hopkins, J., Krietemeyer, J. & Toney, L. Using self-report assessment methods to explore facets of mindfulness. Assessment 13, 27–45 (2006).

Bazin, P.-L. et al. A computational framework for ultra-high resolution cortical segmentation at 7Tesla. Neuroimage 93, 201–209 (2014).

Beck, A. T., Ward, C. H., Mendelson, M., Mock, J. & Erbaugh, J. An inventory for measuring depression. Archives of general psychiatry 4, 561–571 (1961).

Beck, A. T., Steer, R. A. & Brown, G. K. Beck depression inventory-II. San Antonio 78, 490–498 (1996)

Behzadi, Y., Restom, K., Liau, J. & Liu, T. T. A component based noise correction method (CompCor) for BOLD and perfusion based fMRI. Neuroimage 37, 90–101 (2007).

Bertrams, A. & Dickhäuser, O. Messung dispositioneller Selbstkontroll-Kapazität: Eine deutsche Adaptation der Kurzform der Self-Control Scale (SCS-KD). Diagnostica 55, 2–10 (2009).

Bloch, K. E., Schoch, O. D., Zhang, J. N. & Russi, E. W. German version of the Epworth sleepiness scale. Respiration 66, 440–447 (1999).

Carriere, J. S., Seli, P. & Smilek, D. Wandering in both mind and body: individual differences in mind wandering and inattention predict fidgeting. Canadian Journal of Experimental Psychology/Revue canadienne de psychologie expérimentale 67, 19 (2013).

Carson, S. H., Peterson, J. B. & Higgins, D. M. Reliability, validity, and factor structure of the creative achievement questionnaire. Creativity Research Journal 17, 37–50 (2005)

Carver, C. S. & White, T. L. Behavioral inhibition, behavioral activation, and affective responses to impending reward and punishment: The BIS/BAS Scales. Journal of personality and social psychology 67, 319 (1994).

Chang, H. & Fitzpatrick, J. M. A technique for accurate magnetic resonance imaging in the presence of field inhomogeneities. IEEE transactions on medical imaging 11, 319–329 (1992).

Costa, P. T. & McCrae, R. R. The NEO personality inventory manual. (Psychological Assessment Resources., 1985).

Costa, P. T. & McCrae, R. R. Revised NEO personality inventory (NEO PI-R) and NEP five-factor inventory (NEO-FFI): professional manual. (Psychological Assessment Resources Lutz, FL, 1992).

Crowne, D. P. & Marlowe, D. A new scale of social desirability independent of psychopathology. Journal of consulting psychology 24, 349 (1960).

Dale, A. M., Fischl, B. & Sereno, M. I. Cortical surface-based analysis: I. Segmentation and surface reconstruction. Neuroimage 9, 179–194 (1999).

Derryberry, D. & Reed, M. A. Anxiety-related attentional biases and their regulation by attentional control. Journal of abnormal psychology 111, 225 (2002)

Eagleman, D. M., Kagan, A. D., Nelson, S. S., Sagaram, D. & Sarma, A. K. A standardized test battery for the study of synesthesia. Journal of neuroscience methods 159, 139–145 (2007).

Ellison, N. B., Steinfield, C. & Lampe, C. The benefits of Facebook “friends”: Social capital and college students’ use of online social network sites. Journal of Computer-Mediated Communication 12, 1143–1168 (2007).

Esteban, O., Birman, D., Schaer, M., Koyejo, O. O., Poldrack, R. A., & Gorgolewski, K. J. MRIQC: Predicting quality in manual MRI assessment protocols using no-reference image quality measures. bioRxiv 111294 (2017).

Farmer, R. & Sundberg, N. D. Boredom proneness—the development and correlates of a new scale. Journal of personality assessment 50, 4–17 (1986).

Feinberg, D. A. et al. Multiplexed echo planar imaging for sub-second whole brain FMRI and fast diffusion imaging. PloS one 5, e15710 (2010).

Fischl, B., Sereno, M. I. & Dale, A. M. Cortical surface-based analysis: II: inflation, flattening, and a surface-based coordinate system. Neuroimage 9, 195–207 (1999).

Floridou, G. A., Williamson, V. J., Stewart, L. & Müllensiefen, D. The Involuntary Musical Imagery Scale (IMIS). Psychomusicology: Music, Mind, and Brain 25, 28 (2015).

Golchert, J. et al. Individual variation in intentionality in the mind-wandering state is reflected in the integration of the default-mode, fronto-parietal, and limbic networks. NeuroImage 146, 226–235 (2017).

Gorgolewski, K. et al. Nipype: a flexible, lightweight and extensible neuroimaging data processing framework in python. Frontiers in neuroinformatics 5, 13 (2011).

Gorgolewski, K. J. et al. A correspondence between individual differences in the brain’s intrinsic functional architecture and the content and form of self-generated thoughts. PloS one 9, e97176 (2014).

Gorgolewski, K. J. et al. The brain imaging data structure, a format for organizing and describing outputs of neuroimaging experiments. Scientific Data 3, 160044 (2016a).

Gorgolewski, K. J. et al. BIDS Apps: Improving ease of use, accessibility and reproducibility of neuroimaging data analysis methods. bioRxiv, 079145 (2016b).

Gray, J. A. in A model for personality, 246–276 (Springer, 1981).

Gray, J. A. Precis of the neuropsychology of anxiety: An enquiry into the functions of the septo-hippocampal system. Behavioral and Brain Sciences 5, 469–534 (1982).

Greve, D. N. & Fischl, B. Accurate and robust brain image alignment using boundary-based registration. Neuroimage 48, 63–72 (2009).

Guilford, J., Christensen, P., Merrifield, P. & Wilson, R. Alternate uses: Manual of instructions and interpretation. Orange, CA: Sheridan Psychological Services (1978).

Hautzinger, M., Bailer, M., Worall, H. & Keller, F. BDI: Beck-Depressions-Inventar, Testhandbuch, 2. überarbeitete Auflage. Bern: Verlag Hans Huber (1995)

Hawks, R. L. & Chiang, C. N. Urine testing for drugs of abuse. (National Institute on Drug Abuse Rockville, MD, 1986).

Herrmann-Lingen, C., Buss, U. & Snaith, P. Hospital Anxiety and Depression Scale-Deutsche Version (HADS-D). (Huber, 1995)

Hildebrandt, L. K., McCall, C., Engen, H. G., & Singer, T. Cognitive flexibility, heart rate variability, and resilience predict fine-grained regulation of arousal during prolonged threat. Psychophysiology 53(6), 880–890 (2016).

Holmes, A. J. et al. Brain Genomics Superstruct Project initial data release with structural, functional, and behavioral measures. Scientific data 2 (2015).

Hopko, D. R., Mahadevan, R., Bare, R. L. & Hunt, M. K. The abbreviated math anxiety scale (AMAS) construction, validity, and reliability. Assessment 10, 178–182 (2003).

Huettel, S. A. & McCarthy, G. What is odd in the oddball task?: Prefrontal cortex is activated by dynamic changes in response strategy. Neuropsychologia 42, 379–386 (2004).

Jankowska, D. M. & Karwowski, M. Measuring creative imagery abilities. Frontiers in psychology 6, 1591 (2015).

Jenkinson, M. & Smith, S. A global optimisation method for robust affine registration of brain images. Medical image analysis 5, 143–156 (2001).

Jenkinson, M., Bannister, P., Brady, M. & Smith, S. Improved optimization for the robust and accurate linear registration and motion correction of brain images. Neuroimage 17, 825–841 (2002).

Jenkinson, M., Beckmann, C. F., Behrens, T. E., Woolrich, M. W. & Smith, S. M. Fsl. Neuroimage 62, 782–790 (2012).

Jezzard, P. Correction of geometric distortion in fMRI data. Neuroimage 62, 648–651 (2012).

Jezzard, P. & Balaban, R. S. Correction for geometric distortion in echo planar images from B0 field variations. Magnetic resonance in medicine 34, 65–73 (1995).

Joel, D., Tarrasch, R., Berman, Z., Mukamel, M. & Ziv, E. Queering gender: studying gender identity in ‘normative’ individuals. Psychology & Sexuality 5, 291–321 (2014).

Johns, M. W. A new method for measuring daytime sleepiness: the Epworth sleepiness scale. sleep 14, 540–545 (1991).

Jones, D. N. & Paulhus, D. L. Introducing the short dark triad (SD3) a brief measure of dark personality traits. Assessment 21, 28–41 (2014).

Kelly, C., Biswal, B. B., Craddock, R. C., Castellanos, F. X. & Milham, M. P. Characterizing variation in the functional connectome: promise and pitfalls. Trends in cognitive sciences 16, 181–188 (2012).

Kuhl, J. & Kazén, M. Persönlichkeits-Stil-und Störungs-Inventar: PSSI; Manual. (Hogrefe, 2009).

Landmann, N. et al. Entwicklung von 130 deutschsprachigen Compound Remote Associate (CRA)-Worträtseln zur Untersuchung kreativer Prozesse im deutschen Sprachraum. Psychologische Rundschau (2014).

Lee, C. S., Huggins, A. C. & Therriault, D. J. A measure of creativity or intelligence? Examining internal and external structure validity evidence of the Remote Associates Test. Psychology of Aesthetics, Creativity, and the Arts 8, 446 (2014).

Lorio, S. et al. Neurobiological origin of spurious brain morphological changes: A quantitative MRI study. Human brain mapping 37, 1801–1815 (2016).

Lynam, D. R., Smith, G. T., Whiteside, S. P. & Cyders, M. A. The UPPS-P: Assessing five personality pathways to impulsive behavior. West Lafayette, IN: Purdue University (2006).

Marcus, D. S., Olsen, T. R., Ramaratnam, M. & Buckner, R. L. The extensible neuroimaging archive toolkit. Neuroinformatics 5, 11–33 (2007).

Marques, J. P. et al. MP2RAGE, a self bias-field corrected sequence for improved segmentation and T 1-mapping at high field. Neuroimage 49, 1271–1281 (2010).

Mathôt, S., Schreij, D. & Theeuwes, J. OpenSesame: An open-source, graphical experiment builder for the social sciences. Behavior research methods 44, 314–324 (2012).

McCarthy-Jones, S. & Fernyhough, C. The varieties of inner speech: links between quality of inner speech and psychopathological variables in a sample of young adults. Consciousness and cognition 20, 1586–1593 (2011).

Miller, L. C., Murphy, R. & Buss, A. H. Consciousness of body: Private and public. Journal of Personality and Social Psychology 41, 397 (1981).

Moeller, S. et al. Multiband multislice GE-EPI at 7 tesla, with 16-fold acceleration using partial parallel imaging with application to high spatial and temporal whole-brain fMRI. Magnetic Resonance in Medicine 63, 1144–1153 (2010).

Müllensiefen, D., Gingras, B., Musil, J. & Stewart, L. Measuring the facets of musicality: The Goldsmiths Musical Sophistication Index (Gold-MSI). Personality and Individual Differences 60, S35 (2014).

Nooner, K. B. et al. The NKI-Rockland sample: a model for accelerating the pace of discovery science in psychiatry. Frontiers in neuroscience 6, 152 (2012).

O’malley, P. M. & Bachman, J. G. Self-esteem and education: Sex and cohort comparisons among high school seniors. Journal of Personality and Social Psychology 37, 1153 (1979).

Ophir, E., Nass, C. & Wagner, A. D. Cognitive control in media multitaskers. Proceedings of the National Academy of Sciences 106, 15583–15587 (2009).

Ostendorf, F. & Angleitner, A. NEO-Persönlichkeitsinventar (revidierte Form, NEO-PI-R) nach Paul T. Costa und Robert R. McCrae. (Hogrefe, 2004).

Paulhus, D. L. Dark triad of personality (D3-short). Measurement instrument database for the social sciences (2013).

Peirce, J. W. PsychoPy—psychophysics software in Python. Journal of neuroscience methods 162, 8–13 (2007).

Peirce, J. W. Generating stimuli for neuroscience using PsychoPy. Frontiers in neuroinformatics 2 (2008).

Power, J. D. et al. Methods to detect, characterize, and remove motion artifact in resting state fMRI. Neuroimage 84, 320–341 (2014).

Reber, P. J., Wong, E. C., Buxton, R. B., & Frank, L. R. Correction of off resonance-related distortion in echo-planar imaging using EPI-based field maps. Magnetic Resonance in Medicine 39(2), 328–330 (1998).

Rokem, A., Trumpis, M. & Perez, F. in Proceedings of the 8th Python in Science Conference. 68–75 (2009).

Ruby, F. J., Smallwood, J., Engen, H. & Singer, T. How self-generated thought shapes mood—the relation between mind-wandering and mood depends on the socio-temporal content of thoughts. PloS one 8, e77554 (2013).

Sadeghi, H., Hajloo, N., Babayi, K. & Shahri, M. The relationship between metacognition and obsessive beliefs, and procrastination in students of Tabriz and Mohaghegh Ardabili Universities, Iran. Iranian journal of psychiatry and behavioral sciences 8, 42 (2014).

Schaal, N. K., Bauer, A. K. R., & Müllensiefen, D. Der Gold-MSI: replikation und validierung eines fragebogeninstrumentes zur messung musikalischer erfahrenheit anhand einer deutschen stichprobe. Musicae Scientiae 18(4), 423–447 (2014).

Shalev, L., Ben-Simon, A., Mevorach, C., Cohen, Y. & Tsal, Y. Conjunctive Continuous Performance Task (CCPT)—A pure measure of sustained attention. Neuropsychologia 49, 2584–2591 (2011).

Silvia, P. J. et al. Assessing creativity with divergent thinking tasks: Exploring the reliability and validity of new subjective scoring methods. Psychology of Aesthetics, Creativity, and the Arts 2, 68 (2008)

Smith, S. M. et al. A positive-negative mode of population covariation links brain connectivity, demographics and behavior. Nature neuroscience 18, 1565–1567 (2015).

Stöber, J. Tuckman procrastination scale-Deutsch (TPS-D). Unveröffentlichtes Manuskript (1995).

Stöber, J. Die Soziale-Erwünschtheits-Skala-17 (SES-17): Entwicklung und erste Befunde zu Reliabilität und Validität. Diagnostica 45, 173–177 (1999).

Strobel, A., Beauducel, A., Debener, S. & Brocke, B. Eine deutschsprachige Version des BIS/BAS-Fragebogens von Carver und White. Zeitschrift für Differentielle und diagnostische Psychologie (2001).

Tangney, J. P., Baumeister, R. F. & Boone, A. L. High self-control predicts good adjustment, less pathology, better grades, and interpersonal success. Journal of personality 72, 271–324 (2004).

Tuckman, B. W. The development and concurrent validity of the procrastination scale. Educational and psychological measurement 51, 473–480 (1991).

Vaidya, C. J., & Gordon, E. M. Phenotypic Variability in Resting-State Functional Connectivity: Current Status. Brain Connectivity 3, 99–120 (2013).

Van Essen, D. C. et al. The WU-Minn human connectome project: an overview. Neuroimage 80, 62–79 (2013).

Wells, A. & Cartwright-Hatton, S. A short form of the metacognitions questionnaire: properties of the MCQ-30. Behaviour research and therapy 42, 385–396 (2004)

Whiteside, S. P. & Lynam, D. R. The five factor model and impulsivity: Using a structural model of personality to understand impulsivity. Personality and individual differences 30, 669–689 (2001).

Whitmer, A. J. & Banich, M. T. Inhibition versus switching deficits in different forms of rumination. Psychological science 18, 546–553 (2007).

Wittchen, H.-U., Kessler, R. C., Zhao, S. & Abelson, J. Reliability and clinical validity of UM-CIDI DSM-III-R generalized anxiety disorder. Journal of Psychiatric Research 29, 95–110 (1995).

Young, K. S. Internet addiction: The emergence of a new clinical disorder. Cyberpsychology & behavior 1(3), 237–244 (1998).

Zhang, Y., Brady, M. & Smith, S. Segmentation of brain MR images through a hidden Markov random field model and the expectation-maximization algorithm. IEEE transactions on medical imaging 20, 45–57 (2001).

Zigmond, A. S. & Snaith, R. P. The hospital anxiety and depression scale. Acta psychiatrica scandinavica 67, 361–370 (1983).

